# FishWIO: a labeled image dataset of Western Indian Ocean reef fishes for training and testing classification algorithms

**DOI:** 10.64898/2026.03.03.709272

**Authors:** Valentine Fleuré, Thomas Claverie, Sébastien Villéger

**Affiliations:** MARBEC, Univ Montpellier, CNRS, Ifremer, IRD, Montpellier, France; ZooParc de Beauval & Beauval Nature, Saint-Aignan, France; EMR SantEco & UMR ENTROPIE, IRD, IFREMER, CNRS, Univ La Réunion, Saint Denis, 97744, Réunion, France

## Abstract

Monitoring fish communities is essential for understanding biodiversity dynamics and coral reef ecosystem health. Underwater imaging provides a non-invasive and repeatable approach for such monitoring, yet analysis of large volumes of video data remains extremely time-consuming for experts. Resolving such a bottleneck is today within reach, yet towards automated fish identification, large and high-quality, labelled image datasets are critical for training and testing reliable deep learning models. However, to date, no such dataset exists for the Western Indian Ocean (WIO), a global biodiversity hotspot hosting more than 300 common non-cryptobenthic fish species and facing increasing anthropogenic pressures.

This paper presents a novel and publicly available dataset of 114,664 images annotated from 186 videos recorded using fixed underwater cameras on shallow reef habitats from Mayotte archipelago. All images were labelled and validated by trained marine biologists following a standardized protocol. Each image includes detailed metadata describing recording conditions. The dataset comprises 124 reef fish species (including 110 with >200 images) and 8 background classes. This dataset will allow training and testing automated fish classification models.

## Background

Underwater imaging has become a transformative tool for monitoring marine biodiversity (Besson et al., 2022), and has been increasingly used to survey fishes in reef ecosystems for the past decade (Mallet & Pelletier, 2014; Vergés et al., 2016; Dunkley et al., 2023; Loiseau et al., 2023; Magneville et al., 2023). However, the main bottleneck of camera-based censuses is the time required to analyze the recorded images or videos that could exceed the recording time by a two to ten-fold factor (Weinstein, 2018). Automated image analysis to detect and identify fishes has been advancing for the last decade, following the progress in Deep Learning algorithms such as CNN models (Villon et al., 2020; Ditria et al., 2021; Lopez-Marcano et al., 2021; Fleuré et al., 2024; Bürgi et al., 2025). Recent advances in the mathematical basis of Deep Learning such as transformer models pave the way to model able to identify more species with higher efficiency. However, maximizing the performance and generalizability of deep learning models required access to large, taxonomically diverse, and ecologically representative sets of labeled images for training and testing the algorithms (Francescangeli et al., 2023; Contini et al., 2025). Hence, collecting numerous and diverse underwater images and accurately annotating fish individuals at the species level is currently the main bottleneck towards automated monitoring of fish biodiversity (Marrable et al., 2022).

Coral reef fishes are one of the most diverse vertebrate fauna with more than 100 families and 4,000 marine species only in the Indo-Pacific biome (Parravicini et al., 2013). Coral reef fishes are increasingly threatened by the synergistic impacts of climate change, fisheries and pollution (Graham et al., 2011). Even if fishes are among the most studied taxa in the coral reef ecosystems (Brandl et al., 2019), their monitoring through underwater visual censuses (e.g. Edgar & Stuart-Smith, 2014) or underwater videos (e.g. Maslin et al., 2021; Magneville et al., 2024) is demanding for experts. Hence, there is an urgent need for Deep Learning algorithms able to census fish on underwater images. Several open-access image datasets of reef fishes have been released for the last decade (Table 1). However, many of these datasets present limitations that restrict their applicability to monitor biodiversity. For example, the OzFish dataset (Australian Institute Of Marine Science, 2020), contains around 80,000 images for over 500 labels, resulting in a low average number of images per label. In practice, only 12 species were used in the classifier by Marrable (Marrable et al., 2022). Similarly, the dataset from Wang et al. (2025) was gathered from only 23 short videos, averaging 21 seconds each. In addition, videos were all with the same conditions (“clear water, no current, many fishes”), leading to low diversity in backgrounds and lighting conditions. To date, there is no image dataset of reef fishes from the Western Indian Ocean (WIO), a global biodiversity hotspot with more than 300 common non-cryptobenthic species (Samoilys et al., 2022) facing elevated anthropogenic pressures (Poti et al., 2022).

**Table 1:** Overview of published image datasets of reef fishes.

| **Dataset name** | **Reference** | **Region** | **Number of species** | **Number of images** |
| --- | --- | --- | --- | --- |
| Croatian Fish | Jäger et al., (2015) | Adriatic Sea | 12 | 794 |
| Fish4Knowledge | Fisher et al., (2016) | West Pacific Ocean (Taiwan) | 23 | 27,370 |
| WildFish | Zhuang, Wang & Qiao, (2018) | Global | 1000 | 54,459 |
| OzFish | Australian Institute Of Marine Science, (2020) | Eastern Indian Ocean (North West Australia) | 507 | 45,000 |
| DeepFish | Saleh et al., (2020) | West Pacific Ocean (North East Australia) | 2 | 39,766 |
| SeaMap | Boulais et al., (2021) | Western Atlantic Ocean (Gulf of Mexico) | 130 | 90,000 |
| Annotated videos of luderick from estuaries in southeast Queensland, Australia | Ditria et al., (2021) | West Pacific Ocean (East Australia) | 2 | 9,429 |
| FathomNet | Katija et al., (2022) | Global | Not specified | 209,132 |
| Labelled mediterranean fish dataset | Catalán et al., (2023) | Mediterranean Sea (Mallorca Island) | 20 | 17,965 |
| NA | Bürgi et al., (2025) | Mediterranean Sea (French Riviera) | 19 | 68,573 |
| SCSFish2025 | Wang et al., (2025) | West Pacific Ocean (South | 30 | 120,084 |
|  |  | China sea) |  |  |

While citizen-science and collaborative imagery efforts like iNaturalist (https://www.inaturalist.org/) have contributed to the collection of fish images from WIO, they do not specifically aim to publish an image dataset. Seatizen (Contini et al., 2025) provides numerous images of WIO fishes, although they are not annotated at the species level, thus limiting their use for training and testing classification algorithms.

This paper presents a novel and publicly available dataset of 114,664 images of 124 WIO reef fish species and 8 background labels, annotated and validated by experts.

## Data collection

### Image recording

All fish images were extracted from 186 underwater videos recorded between 2015 and 2021, in shallow (0-33m depth) fringing and barrier reefs of Mayotte island (Western Indian Ocean) (Table 2). All videos were recorded stationary near the seabed (20-35cm above seafloor) over various seascapes (i.e. varying relative cover of sand, algae, corals; Figure 1). All videos were recorded using GoPro cameras (Hero 4 Black and Hero 5 Black models) encased in waterproof housings with field-of-view set to ‘linear’ for 95% of them (else ‘wide’), at a resolution of 1920 x 1080 pixels (with 24 to 30 frames per second) and with automatic colour balance. All videos were recorded without any external illumination and thus videos had varying lighting depending on time of the day, cloudiness, depth and water turbidity (Figure 2). The duration of each video varied from less than 1 minute to 17 minutes.

**Table 2:** Summary of image sources. Mayotte hosts two barrier reefs, an inner and outer one, fringing reefs and patch reefs. For clarity, three positions are detailed for the barrier reefs: the outer slope, the reef flat for the top and the inner slope. For example, a site situated on the seaward side of an inner barrier reef is described as “outer slope of inner barrier reef”.

| Campaign date | Sites names | Type of reefs | Number of videos |
| --- | --- | --- | --- |
| November 2015 | Boueni channel;<br>N’Gouja | Inner and outer slope of outer barrier reef; Fringing reef | 49 |
| February 2016 | Submerged barrier reef; N’Gouja; Iris; Iris reef patch | Reef flat and outer slope of outer barrier reef; Fringing reef; Patch reef | 24 |
| March 2016 | Mtsamboro island; Choazil islands; Iris | Fringing reef; Patch reefs | 12 |
| June 2016 | Sakouli; Mtsamboro West; Mtsamboro North; Outer north barrier | Fringing reef; Outer slope of outer barrier reef | 13 |
| September 2016 | Sakouli; Mtsangadoua; Boueni cape | Fringing reef | 11 |
| November 2016 | Chira Rani; Passe Boueni; inner part of the southern barrier; Mronabeja cape; N’Gouja | Fringing reef; Inner and outer slope of outer barrier reef; outer slope of inner barrier reef | 16 |
| Octobre 2017 | Boueni; Barrier; Boueni cape | Fringing reef; outer slope of outer barrier reef | 10 |
| March 2019 | Mwamba harrabou; Vara vara mati; Mwamba maricabou; Gnouma moiba | outer slope of inner barrier reef; Patch reef | 13 |
| November 2020 | Boueni | Fringing reef | 28 |
| March 2021 | Mtsamboro; N’Gouja; Bandrele | Fringing reef | 10 |

**Figure 1:**
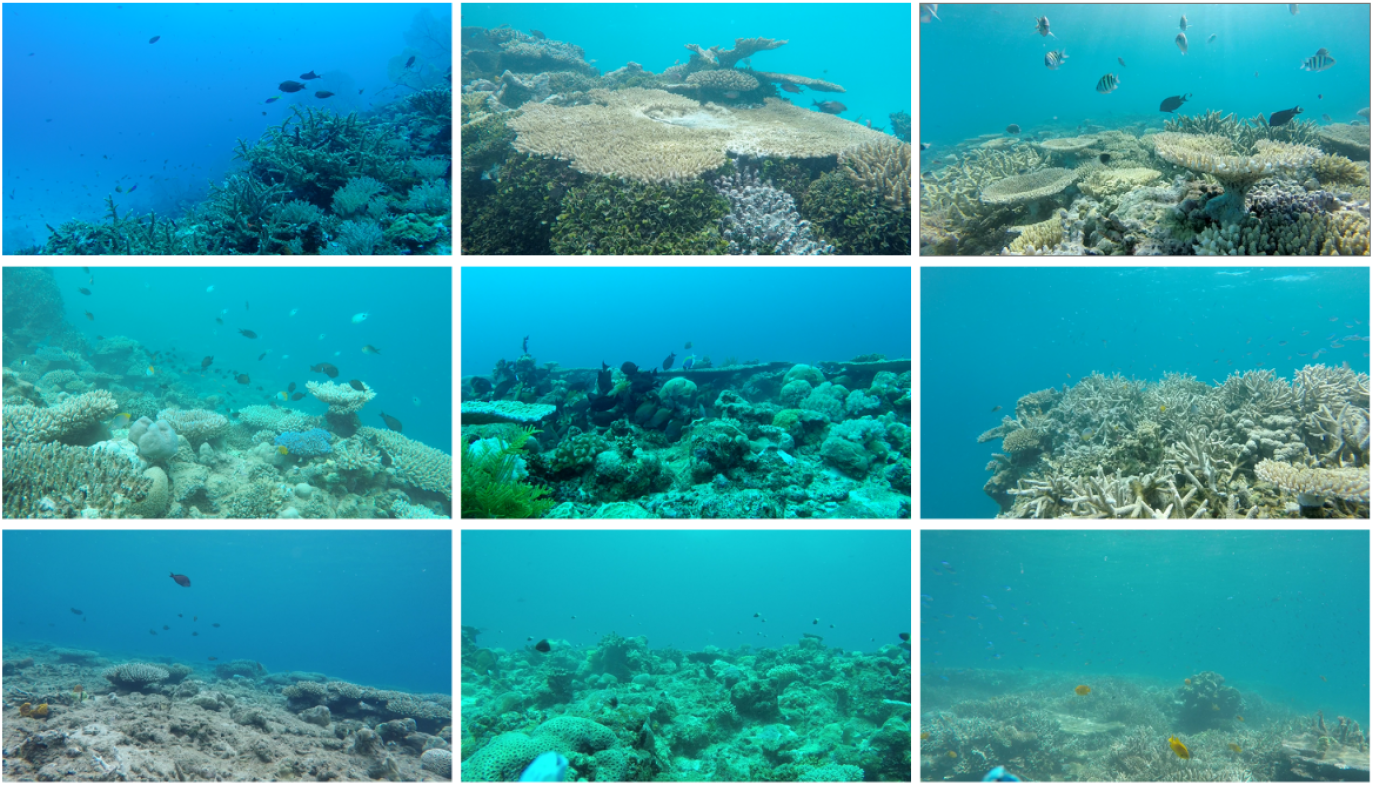
Examples of habitat diversity and environmental conditions present in videos used to build the image dataset.

Videos were eventually grouped into 64 *shootings*, which are defined as recordings captured on the same day and at the same site.

### Fish image labelling

Frames were extracted from the source videos at a rate of 1, 2 or 5 frames per second using the ffmpeg software (https://ffmpeg.org/). These frames were then annotated by more than 30 fish ecologists and supervised bachelor and master students (January 2016 - May 2025).

The protocol was strictly defined and remained consistent for all the annotations. Fish individuals were annotated only if (1) confidently identifiable from the image alone, without the need for contextual cues, surrounding frames, or behavioral clues, (2) >75% body was visible (i.e. not masked by another fish individual or benthos, nor seen from front, back, top or bottom) and (3) fills at least 1,250 pixels (which represent 0.06% of a Full HD frame resolution).

Bounding boxes (i.e. rectangles with sides parallel to video frame sides) were tightly drawn around each fish, minimizing background space and hence aiming to facilitate attention on fish features. Each bounding box was labelled with the species’ latin name (following the GBIF taxonomy; https://www.gbif.org/) and if the species had sexual or ontogenetic dimorphism, the sex (male, female) or life-stage (juvenile, adult) was mentioned as a suffix (e.g. Gomphosus_caeruleus_male). For the species with changing colour patterns due to stress, ambient light or time of the day, we made an effort to annotate all the variants.

Subsequently, for each video, each bounding box was cropped, saved as a jpeg file, and assigned a unique filename consisting of the source video name (including a unique alphanumerical suffix of 13 characters), frame number, width and height in pixel, as well as the pixel coordinates (x, y) of the top-left corner of its bounding box (ex: GOPR0114_592dd43048dc8_250_65x43+1065+187.jpeg).

### Fish images checking

In 2025, experts conducted final validation of all annotations of each species to maintain a uniform standard of quality (i.e. removing erroneously labelled images, images of fish not confidently identifiable, mostly blurred). Taxonomy was checked according to GBIF.org in January 2026.

In cases where more than 200 images of the same species were annotated from a single recording, we randomly subsampled 200 annotations to limit redundancy and reduce sampling bias toward overrepresented seascape or light conditions.

### Additional background images

To further improve the accuracy of classification in a two-stage pipeline (detection followed by classification), the dataset was supplemented with annotation of eight background labels: hard coral, soft coral, calcareous algae, macroalgae, dark background (e.g. shadow under tabular corals, small caves), sand, water column (i.e. blue background), and water surface (i.e. mirror like effects visible). Each background label consists of 200 bounding boxes drawn without any fish on 6 to 16 videos.

## Data records

Images from a specific label were gathered into a unique folder and all label folders were compressed into a single zip file available online (https://doi.org/10.5281/zenodo.17297730).

For each fish image, 14 metadata variables describing the image source and the taxonomy of the fish (Table 3) are available (as a csv file online: https://doi.org/10.5281/zenodo.17297730).

**Table 3:** Description of the metadata collected for each image.

| Column | Description |
| --- | --- |
| Image name | Unique identifier of the image |
| GBIF taxon ID | ID number of the species |
| Order | Order of the species |
| Family | Family of the species |
| Genus | Genus of the species |
| Species | Name of the species |
| Image size | Size of the image in pixels |
| File name | Unique identifier of the video file from which the image was extracted |
| Video date | Date of video recording (YYYYMMDD) |
| Site name | Local name of the site where the videos were recorded |
| Latitude | Latitude of the site where the videos were recorded (SXX.XXXX, where X is a digit) |
| Longitude | Longitude of the site where the videos were recorded (EXXX.XXXX, where X is a digit) |
| Shooting ID | ID of the recording based on the date and geographical coordinates (WGS84; NB: a recording could include several video files) |
| Depth | Depth (rounded to nearest meter) at which the video was recorded |

## Data overview

The final dataset includes a total of 114,664 images, grouped into 158 labels: 150 labels, representing 124 fish species from 25 families and 67 genus, and 8 labels for background categories with 200 images each. Among the 124 fish species, 20 were represented by 2 age stage or sex-based labels and 2 by 3 labels.

There are 753 images on average per fish label (standard deviation = 818), with a median of 455. 32 labels have less than 200 images, 48 labels have between 200 and 500 images, 36 labels have between 500 and 1000 images, and 34 labels have more than 1000 images.

The average size of annotated bounding boxes is 20,210 pixels (1% of a Full HD video frame), with a standard deviation of 26,928 px and a median of 12,540 px. There are 3,793 images between 1250 and 2500 px (0.06%-.1% of Full HD resolution), 11,483 images between 2500 px and 5000 px, 31,229 images between 5000 and 10,000 px, and 68,159 images over 10,000 px. Even small species (i.e. adult body length <15cm) have large images (e.g. 53% of images of *Chromis weberi* >10,000 pixels; Figure 3). On the other hand, large species (i.e. common size >25cm) do have some small images (e.g. 2% of *Naso elegans*) <5,000 pixels.

**Figure 3.**
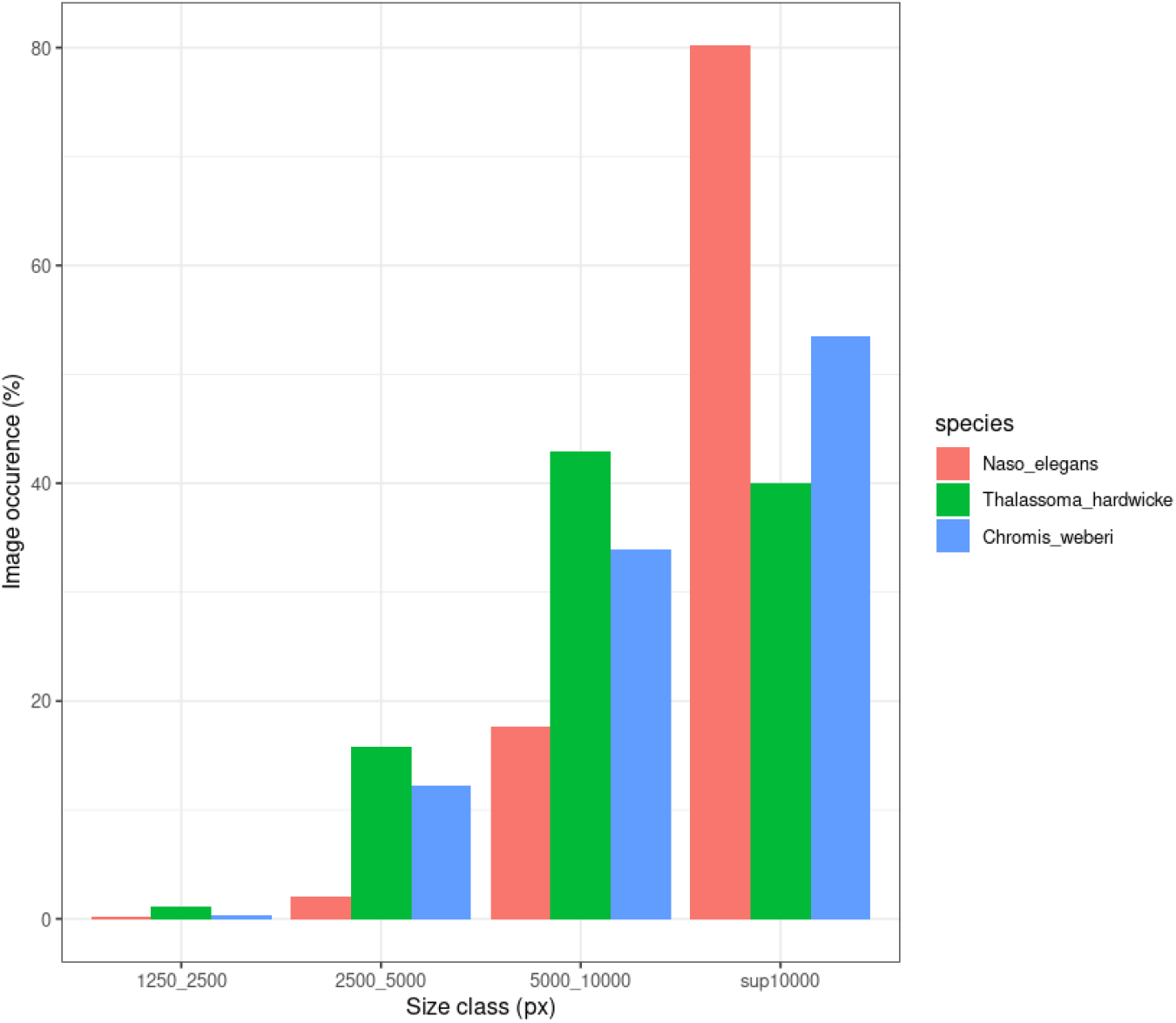
Distribution of image size for three fish species with contrasting common adult body length. The average total length of *Naso elegans* is 45 cm, *Thalassoma hardwicke* is 20 cm, and *Chromis weberi* is 12 cm, size are from (Taquet & Diringer, 2012).

Variability of size is also high among images from each label, with standard deviation being higher than 2000 pixels for all but 2 labels (sd = 982 for Dascyllus_trimaculatus_juv, and sd = 1626 for Pseudodax_moluccanus_adu).

This diversity of image size is favored by our recording protocol. Indeed, as videos were recorded stationary and as in most videos horizontal visibility was >10m the annotated individuals were at markedly variable distance to the camera, hence filling varying areas of video frames (from 1,000 to >10,000 pixels).

In addition, we noted during image checking that the images of a species display a diverse range of aspect ratios, from horizontally to vertically elongated (Figure 4).

**Figure 4:**
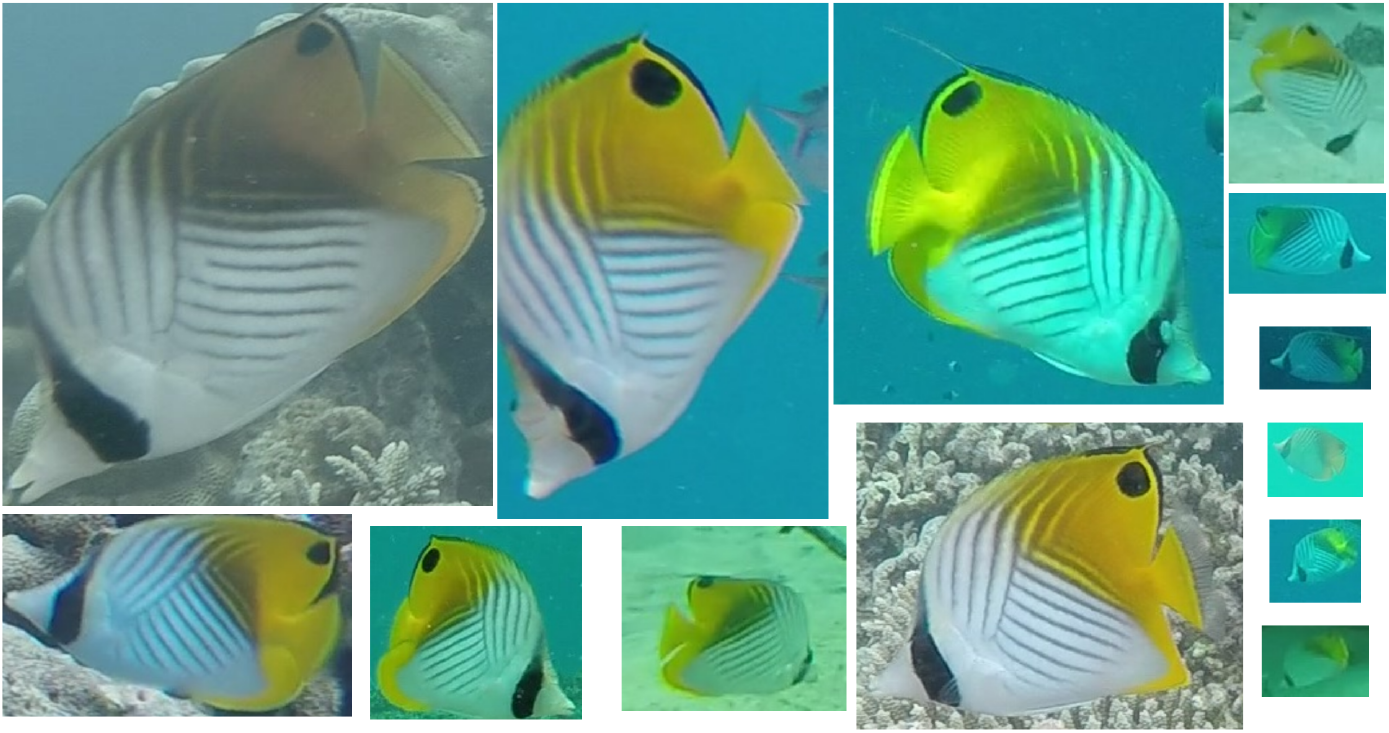
Examples of images from a single label (*Chaetodon auriga*) illustrating variability in fish posture, background and light conditions. The differences in image size are representative of differences found in the dataset; the top left image is 128,800 pixels (6.2% of the frame surface), and the bottom right image is 3,942 pixels (0.19% of the frame surface).

Indeed, the alignment of the fish body with the horizontal axis of the video frame results from their position relative to the camera and their posture that is driven by their movements. Fish often exhibit dynamic orientations during swimming as well as torsion of the whole body or some of its parts (pectoral fins, caudal area). Fish feeding on the seafloor (especially herbivores such as surgeonfishes and parrotfishes) have more often vertical orientation than pelagic fishes feeding on zooplankton (e.g. damselfishes).

## Usage notes

This dataset was assembled to facilitate the training and evaluation of fish species classification models in real-world underwater conditions. The annotation protocol ensures that only individuals that can be reliably identified based on a single frame are included in the dataset. In addition, the inclusion of several labels for species with dimorphism will improve the training efficiency and ultimately the precision of classification. The large number of videos used to extract fish images as well as the annotation protocol favor the diversity of images (size, body orientation, ambient light) within each species. This diversity could even be increased using dataset augmentation techniques such as horizontal flipping and vertical and horizontal slicing that improve training efficiency (Villon et al., 2018). Furthermore, the metadata associated with each image includes the recording and hence eases the split of images of each species into independent train and test datasets (i.e. ensuring tests include only images from videos not used to build the train set).

Convolutional neural network models performed well only with >1,000 images per species which drastically limits the number of species included (e.g. Villon et al. (2020), which was limited to 20 species from WIO). The recent transformer-based models which are efficient even with only 200 images per species (Wang et al., 2025) will allow training classifiers with our dataset for 108 species (including all the ones from Villon et al 2020).

Of the 124 species included in our dataset, 92 are also present in the OzFish dataset (Australian Institute Of Marine Science, 2020) and 16 in Wang et al. (2025). Our dataset therefore contributes 34 species not represented in the OzFish dataset (Australian Institute Of Marine Science, 2020) and 110 species not represented in the SCSfish dataset (Wang et al., 2025).

Moreover, the inclusion of the background labels is useful to reduce the risk of erroneously classifying false positives from the detection stage (Villon et al 2018) or even through training a single-class classifier (Jenrette et al., 2022).

Furthermore, the gathering of this dataset was part of a strategy including a fish detector followed by a species classifier instead of a multiclass detectors. Indeed, according to our protocol, only some individuals matching our quality criteria were annotated on each frame, and this strategy allowed us to quickly collect thumbnails of rare species. Therefore, we acknowledge that our dataset can not be used to train multi-label detection algorithms that require frames with all fishes annotated (even those not identifiable), But, such ready to train fish detector dataset for Indo-Pacific reef fishes are available (Wang et al., 2025).

The objective is to continuously update this dataset and the associated metadata table as data collection progresses. This includes adding more images for species with fewer than 200 images and incorporating new species, including congeneric species that are highly similar to each other (e.g *Forcipiger flavissimus* and *F. longirostris*)

## Data Availability

The dataset is available at https://doi.org/10.5281/zenodo.17297730

## Acknowledgement

This work was funded by the Association Beauval Nature (CIFRE 2022/0127) and by the IA-Biodiv ANR project FishPredict (ANR-21-AAFI-0001-01). The CUFR (now University of Mayotte) funded an important part of the logistics for camera deployment.

We thank Anne-Sophie Tribot, Sébastien Villon, Elliott Sucré, Yann Mercky, Charles Le Bozec, Marie Gimenes, Camille Magneville and Nicolas Loiseau who helped to record the videos over the years.

We thank Clément Desgenetez for designing the annotation tool. We thank the 30 students from the University of Montpellier who helped to annotate fishes on video frames during these years.

